# Biased competition in the absence of input bias: predictions from corticostriatal computation

**DOI:** 10.1101/258053

**Authors:** Salva Ardid, Jason S. Sherfey, Michelle M. McCarthy, Joachim Hass, Benjamin R. Pittman-Polletta, Nancy Kopell

## Abstract

Classical accounts of biased competition (BC) require an input bias to resolve the competition between neuronal ensembles driving downstream processing. However, flexible and reliable selection of behaviorally-relevant ensembles can occur with unbiased stimulation: striatal D1 and D2 spiny projecting neurons (SPNs) receive balanced cortical input, yet their activity determines the choice between GO and NO-GO pathways in the basal ganglia. We present a corticostriatal model identifying three mechanisms that rely on physiological asymmetries to effect rate- and time-coded BC in the presence of balanced inputs. First, tonic input strength determines which SPN phenotype exhibit higher mean firing rate (FR). Second, low strength oscillatory inputs induce higher FR in D2 SPNs but higher coherence between D1 SPNs. Third, high strength inputs oscillating at distinct frequencies preferentially activate D1 or D2 SPN populations. Of these mechanisms, the latter accommodates observed rhythmic activity supporting rule-based decision making in prefrontal cortex.

---

Biasing the competition between neuronal ensembles is essential for preferential processing of relevant visual information (*1*). Two computational principles underlie biased competition, as currently understood. First, stimulus-driven neuronal ensembles having distinct stimulus selectivity suppress each other’s activity via mutual inhibition. Second, an external input preferentially targets one of the competing ensembles, breaking the symmetry of the system. Computational models of biased competition implementing these principles can differ in considering either an asynchronous (*2, 3, 4, 5*) or a rhythmic (*6*) input bias, as well as in the impact of the bias on neural circuit dynamics, which may increase firing rate (FR) (*2, 3*), coherence (*4, 5*), or both (*6*).

In this work, we introduce an entirely different set of computational principles for biased competition. In the absence of externally imposed (i.e. input) biases, we will show that biased competition is possible between neuronal ensembles endowed with distinct physiological properties. We use corticostriatal processing as a model system for biased competition in the absence of an input bias, because striatal input-output processing is mediated by competition between two distinct GABAergic populations of spiny projecting neurons (SPNs), preferentially expressing either D1 or D2 dopamine receptors (*7, 8, 9*), that receive balanced cortical stimulation (*10*).

The manifold differences between D1 and D2 SPNs span anatomical (*11*), network (*12*) and intrinsic properties (*13*), and the two inhibitory populations interact in complex and asymmetrical ways. Thus, it is difficult to predict which physiological asymmetries enable biased competition, and under which conditions. Consequently, we addressed this question in a neural circuit model of corticostriatal processing (Fig. 1A). In this model, D1 and D2 SPNs (which we think of and refer to as ensembles, respectively representing to execute or hold an action) receive balanced cortical stimulation (*10*), and exhibit the three main experimentally observed physiological differences: (i) an asymmetric connectivity profile (Fig. 1A), in which there are about five times more connections from D2 to D1 SPNs, than vice-versa (*11*); (ii) distinct GABAergic dynamics (Fig. 1B), with efferent synapses from D1 SPNs having higher maximum conductance but more rapid depression than those from D2 SPNs (*12*); and (iii) intrinsic properties (Fig. 1C), such that outward calcium-dependent potassium currents are activated earlier and more strongly in D2 SPNs (*13*).

**Fig. 1.**
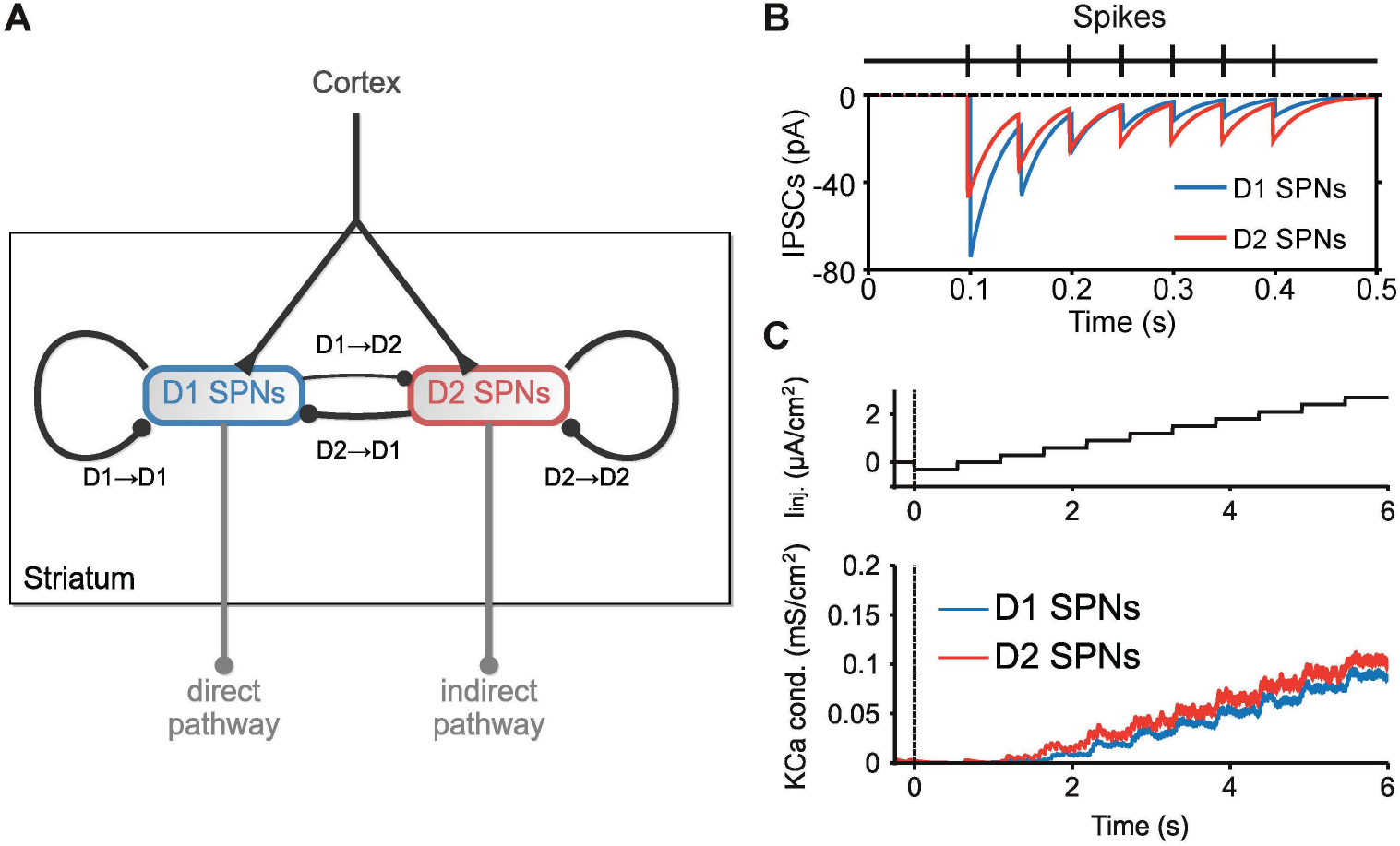
Corticostriatal circuit model. (**A**) The model of the striatum is composed of D1 and D2 spiny projecting neurons (SPNs) according to expressed dopamine receptor. Both phenotypes receive external, balanced input from cortex. D1 and D2 SPNs respectively represent the first stage of the direct (GO) and the indirect (NO-GO) pathways of the basal ganglia. There is an asymmetric connectivity between D1 and D2 SPNs: about 5% of D1 SPNs target D2 SPNs, whereas other synaptic connections in the microcircuit vary within the 20-35% range. (**B**) Distinct GABAergic dynamics between D1 and D2 SPNs. Synapses emerging from D1 SPNs have higher GABAergic conductance (see the difference in amplitude of the first IPSC), but they get depressed more rapidly (see the evolution of IPSC amplitudes). (**C**) Higher activation of outward calcium-dependent potassium currents in D2 SPNs. Top panel shows the protocol of injected current that is applied to D1 and D2 SPNs in the model. Bottom panel shows earlier and stronger activation of the channel for D2 SPNs.

Functionally, D1 and D2 SPNs represent the first relay of the direct (GO) and indirect (NO-GO) pathways of the basal ganglia (Fig. 1A). GO and NO-GO pathways compete with each other to either trigger or hold an action (*14*). Coactivation of D1 and D2 SPNs during action initiation (*15*) imposes a limitation on winner-take-all competition in the striatum (*16*). Recent modeling work proves, however, that even weak activity biases strongly influence downstream attractor dynamics subserving routing and decision making (*17*). Our corticostriatal model is consistent with this view. While the time course of a selected action may depend on complex interactions between basal ganglia nuclei (*16, 18*), a bias in the activity of D1 and D2 SPNs may be sufficient to determine that selection.

But, how can balanced input enable a flexible biasing of neuronal ensembles–i.e., one that allows the reliable selection of either ensemble through variation in the properties of their common input—, in the first place? We hypothesized, and confirmed in our model, two distinct types of mechanisms, each exploiting a specific neural code based on mean firing rate or on precise spike timing (coherence).

First, tonic input strength is able to induce a flexible mean firing rate bias, in which each neuronal ensemble is preferentially activated by inputs within a characteristic range of intensities (Fig. 2C). This mechanism applies even though the two ensembles have the same baseline activity (Fig. 2A and B). The fact that the two ensembles are differentially activated by high and low intensity inputs results from a trade-off between inhibition and activity-dependent hyperpolarization: higher GABAergic inhibition targeting D1 SPNs predominates at low input strengths, leading to higher FR in D2 SPNs, whereas higher outward calcium-dependent potassium currents in D2 SPNs reverse this bias at high input strengths (Fig. 1C). This turning point in relative excitability fits with a confidence-based action selection interpretation of corticostriatal computation: low input strengths represent low signal-to-noise ratios, for which triggering NO-GO actions may be behaviorally advantageous. Thus, the NO-GO pathway is favored at low confidence levels, i.e., when SPNs receive asynchronous low strength inputs, whereas the GO pathway is favored at high confidence levels, i.e., when SPNs receive asynchronous high strength inputs. The mechanism is only apparent at the population level, when a sufficiently large proportion of cells are stimulated by the input. In contrast, the turning point disappears under single-cell stimulation (Fig. 2C, inset), because single-cell stimulation barely affects GABAergic dynamics.

**Fig. 2.**
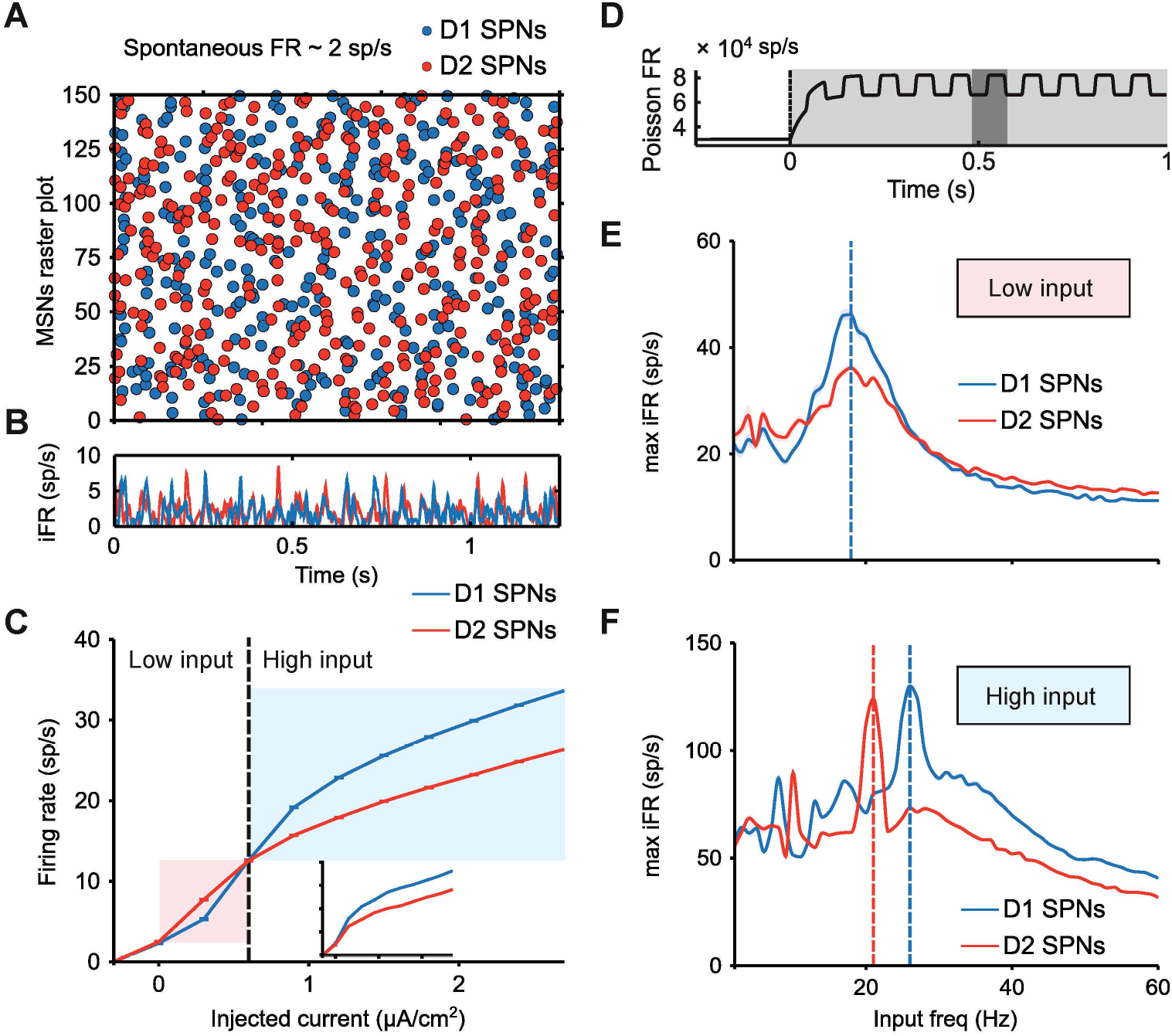
D1 and D2 SPNs respond differently to balanced input. (**A**) Raster plot of the spontaneous activity of SPNs. (**B**) Instantaneous firing rate (iFR, average firing of the population varying in time) of SPNs. (**C**) Averaged f-I curve of each neuronal ensemble (input-output transfer function between injected current, as in Fig. 1C top, and time-averaged population firing rate). The higher excitability of D2 SPNs is shaded in red. The higher excitability of D1 SPNs is shaded in blue. Inset plot: f-I curve when the injected current is applied only to a single cell of each population. (**D**) Poisson rate of the oscillatory input to the striatal circuit: when the stimulus is on (shaded in light gray), the Poisson rate increases and oscillates at a given frequency (inverse of the period, which appears shaded in dark gray). (**E**) Resonance of SPNs for low strength input. The resonance is quantified in terms of maximum iFR, a measure of local population synchronization: across input frequencies (x-axis), the average of the peak iFR through all cycles is computed (y-axis). (**F**) Resonance of SPNs for high strength input. Computed as in (E).

Second, an oscillatory input can induce a coherence bias by preferentially activating the resonant properties of a specific neuronal ensemble. In fact, by varying the frequency of a rhythmic cortical input (Fig. 2D), our model reveals two ways in which the resonances of the two SPN ensembles may diverge: At low input strength, D1 and D2 SPN populations both resonate at the same (low beta) frequency, but D1 SPNs are much more strongly synchronized by rhythmic input (Fig. 2E), despite their lower FR. This divergence between rate and coherence relies on synaptic inhibition. Higher inhibition decreases the overall FR, but enhances spiking coherence, since cells are pushed closer to baseline by inhibition and thus exhibit a more uniform state when inhibition wears off (*19*). At high input strengths, the resonant frequencies of D1 and D2 SPN populations both increase, and diverge from each other (Fig. 2F). The increases in resonant frequency occur because the external input drives SPNs faster than their network frequency in the low beta band (Fig. 2E). As a result, the resonant frequencies of D1 and D2 SPNs shift beyond low beta, respectively to high and middle beta frequencies, following their mean FR (Fig. 2C).

Temporal coordination is, in addition to increased input strength, another mechanism enhancing the signal-to-noise ratio. Thus, while it seems behaviorally advantageous to favor the NO-GO pathway under low strength asynchronous inputs, reliable GO selections can be accomplished for low strength inputs when oscillatory frequency matches the resonant dynamics of SPNs. Furthermore, our results suggest that highly reliable selections of GO and NO-GO actions (in the sense of highest signal-to-noise ratios) occur under rhythmic inputs of high strength, when the oscillatory frequency of the input specifically matches either the resonant frequency of D1 or D2 SPNs.

All together, Figure 2 predicts three types of inputs supporting flexible biased competition under balanced stimulation, confirmed in Figure 3 (top and middle): (i) a rate bias regulated by high vs. low input strength (Fig. 3A and B); (ii) a coherence bias regulated by high strength inputs oscillating at distinct beta bands (Fig. 3D and E); and (iii) coexisting rate and coherence biases in the activity of D2 and D1 SPNs, respectively, resulting from low strength oscillatory inputs at low beta frequency (Fig. 3C). But, how reliable is each bias at driving downstream action selection?

**Fig. 3.**
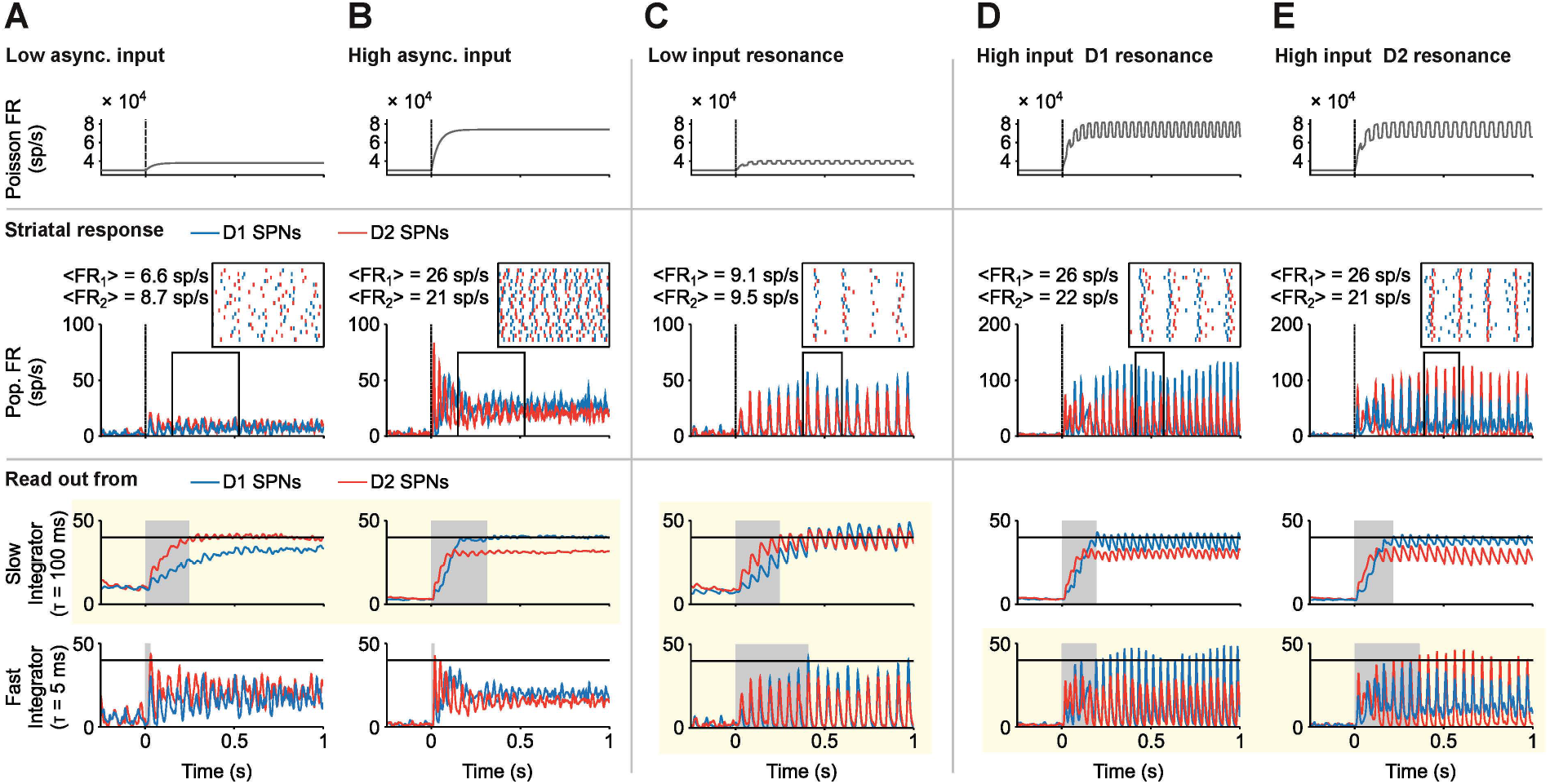
Three successful mechanisms that flexibly bias the striatal circuit under balanced input. (**A** vs. **B**) Biasing between GO vs NO-GO pathways may depend on the overall strength of the balanced cortical input. Top: Poisson rate of the balanced, asynchronous input to the striatal circuit. Middle: Population FR varying in time (iFR). Inset plot: raster activity of (n = 20) D1 and D2 SPNs (time window indicated by the black frame in main plot). Bottom: Downstream readout of the activity of SPNs using distinct integration timescales. Only slow timescale integration shows flexibility in biasing between D1 and D2 SPNs (highlighted in yellow background). Response time is shaded in light gray. Response threshold at 40 sp/s (solid horizontal line). (**C**) Biasing between GO vs NO-GO pathways under balanced inputs of low strength may depend on resonant properties of SPNs and a dynamic tuning of readout timescale. Top: Poisson rate of the balanced, oscillatory input to the striatal circuit that matches the resonant frequency of SPNs (low beta). Middle: Population FR varying in time (iFR, whose amplitude is a measure of local population synchronization). Inset plot: raster activity of (n = 20) D1 and D2 SPNs (time window indicated by the black frame in main plot). Bottom: Downstream readout of the activity of SPNs using distinct integration timescale. On-the-fly tuning of the readout timescale allows flexibility in biasing between D1 and D2 SPNs (highlighted in yellow background). Response time is shaded in light gray. Response threshold at 40 sp/s (solid horizontal line). (**D** vs. **E**) Biasing between GO vs NO-GO pathways under balanced inputs of high strength may depend on resonant properties of SPNs and the spectral content of the balanced cortical input. Top: Poisson rate of the balanced, oscillatory input to the striatal circuit that matches the resonant frequency of either SPN type (high and middle beta, respectively). Middle: Population FR varying in time (iFR, whose amplitude is a measure of local population synchronization; note the higher scale of (D) and (E) compared to (A)-(C)). Inset plot: raster activity of (n = 20) D1 and D2 SPNs (time window indicated by the black frame in main plot). Bottom: Downstream readout of the activity of SPNs using distinct integration timescale. Only fast timescale integration shows flexibility in biasing between D1 and D2 SPNs (highlighted in yellow background). Response time is shaded in light gray. Response threshold at 40 sp/s (solid horizontal line).

To address this question we ran the model output through two read-out decoders of striatal activity. The first decoder was a spiking activity accumulator, with a slow integration timescale (*τ* = 100 ms). The second decoder was a coincidence detector, with a fast integration timescale (*τ* = 5 ms). Our results show that the nature of the striatal bias must fit the timescale of the decoder to guarantee reliable downstream selection (Fig. 3). Thus, only the activity accumulator reliably selects either the GO or the NO-GO pathway from the FR bias between D1 and D2 SPNs (Fig. 3A and B, bottom), while only the coincidence detector flexibly selects either the GO or the NO-GO pathway from the coherence bias between D1 and D2 SPNs (Fig. 3D and E, bottom). When rate and coherence biases coexist, the selection between GO and NO-GO pathways depends on the integration timescale of the decoder. Thus, a coincidence detector reads out the coherence bias of D1 SPNs, whereas an activity accumulator reads out the rate bias of D2 SPNs. Flexible action selection in this case requires adjusting the decoder integration timescale, so it behaves as a coincidence detector or as an activity accumulator. One way to accomplish this may be adjusting the amount of balanced inhibition targeting the decoder, which has been shown to regulate temporal precision (*20*).

These mechanisms impose predictions that can be tested experimentally. According to the rate bias mechanism, action release must functionally correlate with higher mean FR of D1 over D2 SPNs (Fig. 3B). According to the coherence bias mechanism, action release must functionally correlate with a peak in spike-field coherence at either high (Fig. 3D) or low (Fig. 3C) beta frequencies. Note that mean rate and coherence biases may both be present at once (e.g., Fig. 3D). In this context, a contrast between correct vs. error trials, and/or between short vs. long response time trials, may help to identify the ultimate mechanism supporting BC. Thus, in Figure. 3D, we expect that the amplitude of the peak in spike-field coherence is more strongly correlated with behavior than the mean rate difference between D1 and D2 SPNs.

We have focused so far on the situation in which D1 and D2 SPN ensembles compete for the power to trigger or hold isolated actions, but most frequently goal-directed behaviors require selecting the proper action from multiple available sensory-motor associations, such as in rule-based decision tasks. Rule, category and stimulus selective neural activities have been reported in prefrontal cortex (PFC) (*21, 22, 23, 24*) and striatum (*25, 26*), with coactivation of competing ensembles in PFC, coactivating, in turn, competing pathways in the basal ganglia (*27*). Modeling studies have proposed connectionist and rate coding mechanisms to describe routing of sensory-motor responses according to rule biases (*28, 29, 17*); however, recent experimental evidence highlights the central role of temporal dynamics. In particular, rhythmic activity at high beta frequencies is observed in the interaction between PFC and striatum during category learning (*30*), as well as within PFC while performing a rule-based decision task (*31*), where beta phase-locking was higher for the neuronal ensemble encoding relevant information than for its irrelevant competitors. In the same rule-based decision task, alpha-band prefrontal activity was suggested to mediate suppression of ensembles processing the dominant sensory-motor responses during non-dominant trials, i.e., when these representations were irrelevant (*31*).

Our model sheds light on how high beta and alpha rhythms might affect downstream processing in the basal ganglia during this task, suggesting a coherence bias as a mechanistic explanation for rule-based action selection based on stronger high beta synchronization of relevant ensembles in PFC. We hypothesize that while D1 and D2 SPN ensembles representing the same categorical action receive balanced input, SPNs representing relevant categorical actions receive more synchronized input at high beta frequency than SPNs representing irrelevant categorical actions (Fig. 4A top). Higher input synchrony produces more coherent striatal firing (Fig. 4A middle), a bias that can be reliably read-out by a coincidence detector, but not by an activity accumulator because the mean FR is the same for relevant and irrelevant SPN ensembles (Fig. 4A middle and bottom). Thus, higher beta coherence in PFC is able to bias relevant over irrelevant GO pathways of the basal ganglia. Importantly, neither of the other two biased competition mechanisms present in our model favored the relevant GO pathway (Fig. S3B and supplementary text).

**Fig. 4.**
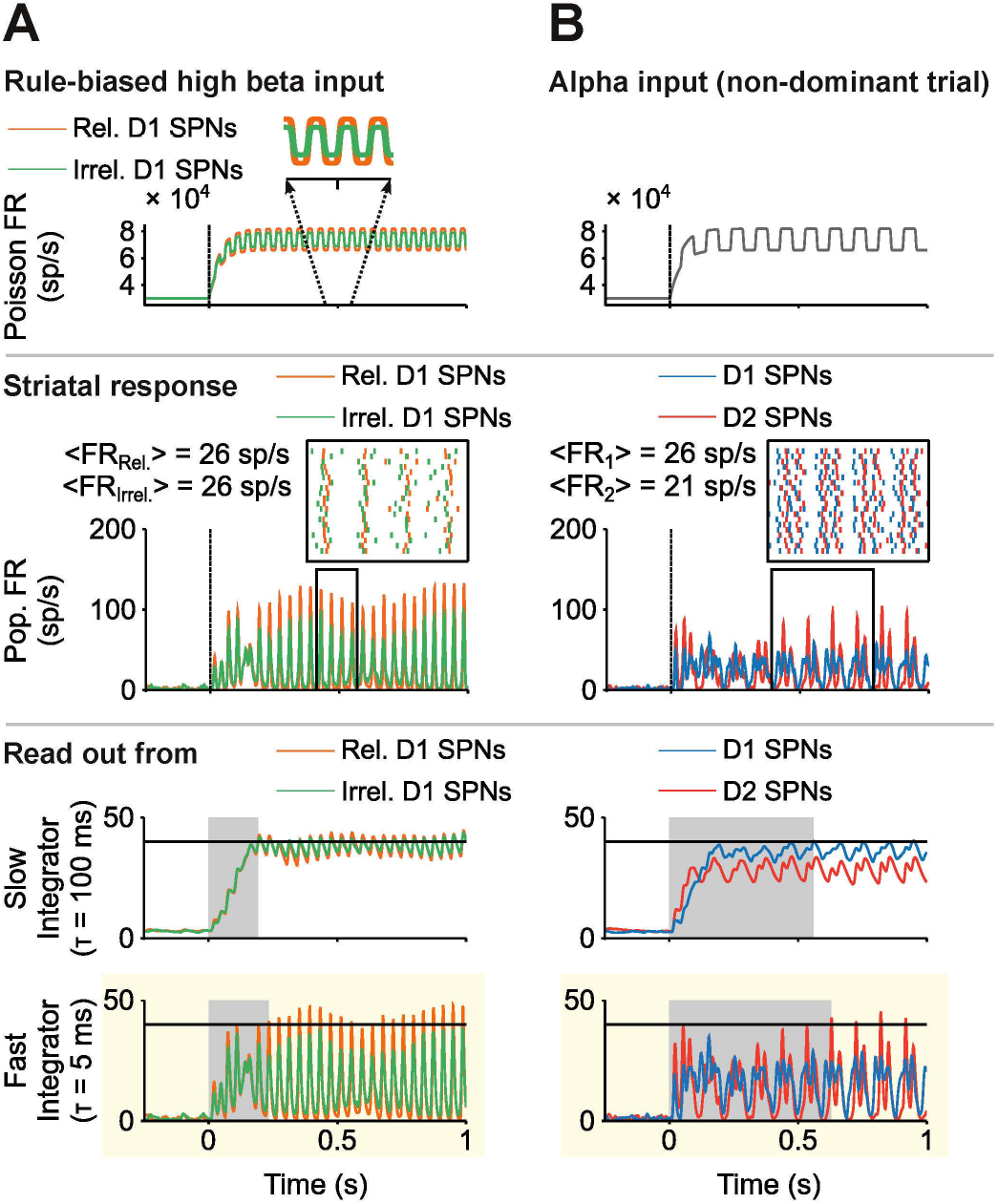
Striatal processing of rhythmic cortical inputs involved in rule-based decisions. (**A**) Rule-based biased competition between GO pathways: stimulus-driven high beta rhythmic input from PFC biases action selection in the basal ganglia. Top: Poisson rate of the high beta oscillatory input to the striatal circuit. The frequency (high beta) and distinct amplitude of the oscillatory input to SPNs (higher for relevant SPNs; see inset plot) are constrained by reported synchronous activity in PFC (see text for details). The input was balanced with respect to D1 and D2 SPNs. Middle: Population FR varying in time (iFR, whose amplitude is a measure of local population synchronization). Inset plot: raster activity of (n = 20) relevant and irrelevant D1 SPNs (time window indicated by the black frame in main plot). Bottom: Downstream readout of the activity of D1 SPNs using distinct integration timescale. Only fast timescale integration shows a reliable bias in favor of the relevant D1 SPN ensemble (highlighted in yellow background). Response time is shaded in light gray. Response threshold at 40 sp/s (solid horizontal line). (**B**) Striatal dynamics toward the NO-GO pathway can be favored by an alpha oscillatory input, present in PFC ensembles in non-dominant trials (see text for details). Top: Poisson rate of the balanced, alpha oscillatory input to the striatal circuit. Middle: Population FR varying in time (iFR). Inset plot: raster activity of (n = 20) D1 and D2 SPNs (time window indicated by the black frame in main plot). Bottom: Downstream readout of the activity of SPNs using distinct integration timescale. For the alpha rhythm to be associated with inhibitory control, D2 SPN bias should prevail over D1 SPN bias. This is only the case through fast integration timescale (highlighted in yellow background). Response time is shaded in light gray. Response threshold at 40 sp/s (solid horizontal line).

In the basal ganglia, inhibitory control is mediated by the indirect (NO-GO) pathway. For an alpha rhythm in PFC to play a role in downstream inhibitory control, it would have to bias the activity of D2 over D1 SPNs. This is the case for the coherence bias mechanism: a balanced cortical input oscillating at alpha frequencies (Fig. 4B top) leads to more coherent firing in D2 SPNs (Fig. 2F and Fig. 4B middle), which can be reliably read-out by a coincidence detector (Fig. 4B bottom). An activity accumulator, on the contrary, does not support an alpha oscillatory input as an inhibitory control mechanism, since it reads out the higher mean FR of D1 SPNs (Fig. 4B bottom). Thus, our model suggests a manner by which cortical inputs oscillating at alpha frequencies synchronize the activity of D2 SPNs more strongly than that of D1 SPNs, hence favoring the selection of the NO-GO pathway. Neither of the other two biased competition mechanisms favored the NO-GO pathway (Fig. S3A and supplementary text).

The results reported in this work reveal novel computational principles underlying preferential processing in support of goal-directed behaviors, such as action selection in the striatum. These mechanisms extend previous approaches that only considered unbalanced inputs as the source of the bias between competing neuronal ensembles. In the context of corticostriatal processing, such an approach (*32*) is challenged by the evidence of balanced cortical input to SPNs (*10*). In contrast, our model predicts, to our knowledge for the first time, that flexibly biasing basal ganglia dynamics toward activation of either the direct or the indirect pathway can be accomplished by tuning either the strength or the spectral properties of a balanced cortical input. Of the alternative mechanisms supporting BC under balanced input, only the coherence bias mechanism is consistent with observed rhythmic activity in PFC in the context of rule-based decisions (*31*). In fact, our model of corticostriatal processing suggests a mechanistic explanation for how alpha and high beta rhythms in PFC support, respectively, inhibitory control and rule-based action selection in the basal ganglia.

An attractive, if speculative, hypothesis is that the three BC mechanisms play a role at different learning stages. Dopamine release increases the firing rate of rule-selective neural ensembles in the PFC (*33*), and these very same ensembles synchronize at high beta frequency (*31*), which is expected to build-up through training. Based on these observations, we suggest that corticostriatal inputs grow in signal-to-noise ratio, both in strength and coherence, through practice. Thus, at early learning stages, cortical inputs are presumably of weak intensity, for which NO-GO activation may be the default mode (Fig. 3A), unless inputs to SPNs temporally coordinate at low beta frequency (Fig. 3C). Later on, through continuous learning, cortical inputs are expected to grow in mean drive, increasing the signal to noise ratio and then biasing the preferential selection toward the GO pathway further and further (Fig. 3B). At this point the task is mastered, so that cortical inputs are strong enough to dissociate the resonant frequency of SPNs (Fig. 3D vs. 3E), so that alpha vs. high beta inputs are able to reliably activate top-down triggered inhibitory control (Fig. 4B) vs. rule-based action selection (Fig. 4A).

The validity of these computational principles may extend beyond corticostriatal processing. Thus, a rate bias may arise wherever a difference in relative excitability exists between competing neuronal ensembles (*34*), and a coherence bias may be induced whenever competing neuronal ensembles resonate at distinct frequencies (*35*). For the two biases to exist simultaneously, there must be a trade-off between FR and coherence. In our model, this trade-off relies on competing neuronal ensembles receiving different amounts of inhibition, internally generated within the striatal microcircuit, despite balanced cortical input. We suspect that the FR-coherence trade-off may also be present when competing ensembles have different AMPA to NMDA conductance ratios, resulting in different synaptic decay timescales: while more AMPA excitation may enhance coherent dynamics (6), less NMDA excitation reduces the overall excitability and, hence, decreases mean FR.

We analyzed biased competition between distinct neuronal ensembles receiving the same inputs, the inverse condition of “classical” biased competition, where identical ensembles receive unbalanced inputs. In general, however, biased competition may occur between competing neuronal ensembles that differ both physiologically and in their input. While this situation is more complex (e.g., cooperation vs. competition between internal and external biased competition mechanisms), it may also be more ubiquitous in the brain and, hence, important to consider systematically. Our work provides a foundation upon which to start addressing this challenge.

## Acknowledgments

This research was supported by ARO Grant W911NF-12-R-0012-02. N.K. and S.A. were also supported by NSF Grant DMS-1042134. M.M.M. was supported by CRCNS NIH Grant CRCNS 1R01NS081716. S.A. designed research; N.K. supervised research; All authors contributed to guide research in regular discussions; S.A. implemented the model, with contributions from M.M.M. and J.S.S.; S.A. ran the simulations, analyzed data, prepared the figures and wrote the manuscript; All authors contributed to edit and revise the text. We thank T. Womels-dorf for helpful suggestions on the manuscript.

## Supplementary Text

There are two main points we focus on in this supplementary section. First, we compare our striatal circuit model with previous striatal modeling work in the lab. And second, we describe how the two mechanisms supporting biased competition under balanced input, alternatives to the coherence bias mechanism, fail to be consistent with observed rhythmic activity in PFC in the context of rule-based decisions.

1. Our striatal circuit model builds upon previous work in the lab (*36*). Our current implementation differs, however, from the previous model in four relevant aspects:

- Two SPN phenotypes: this version of the model considers the two main phenotypes of spiny projecting neurons, D1 and D2 SPNs. D1 and D2 SPNs together constitute about 95% of the cells in the striatum and, unlike other cell types in the striatum, SPNs are the cells receiving the majority of inputs and the only cells projecting downstream to other nuclei of the basal ganglia (*37*). For simplicity, our model bypassed other cell types of the striatal circuit in the model, under the assumption that they do not play a major role in controlling the dynamics of the striatal circuit in the context of preferential corticostriatal processing beyond modulating the overall level of inhibitory input to SPNs. We nevertheless acknowledge the large diversity of interneurons in the striatum. Future computational studies should further analyze the contribution of striatal interneurons in rule-based decisions, and their particular influence, if any, in the striatal response to prefrontal inputs at alpha and beta frequencies. D1 and D2 SPNs possess an asymmetric connectivity profile (*11*) (Fig. 1A). To analyze how the asymmetric, sparse connectivity affects circuit dynamics, we compared the control population-averaged f-I curves (Fig. 2C) with the same f-I curves assuming all-to-all connectivity (Fig. S1). The comparison of f-I curves reveals that no substantial difference is found if the maximal conductance is rescaled by the connectivity ratio. Thus, the main role of asymmetric connectivity in circuit dynamics is to modulate the GABAergic directionality between D1 and D2 SPNs.
- Health condition: our model aims to represent the dynamics of SPNs in health condition, whereas the previous model was a striatal model of Parkinson’s Disease (PD), which assumed dopamine depletion. The overall increase in dopamine concentration in health compared to *in vitro* recordings or PD (*7, 38*) affects the intrinsic excitability of each SPN phenotype in opposite directions (*14*): while D2 SPNs excitability strongly decreases, D1 SPNs excitability increases, up to a point where no apparent differences exist between the baseline activity of D1 and D2 SPNs (*39*). Dopamine concentration ([DA]) also impacts neural dynamics at the circuit level via neuromodulation of GABAergic synapses (*12*) (Fig. 1B). In our model, we considered these high vs. low [DA] *in vitro* recordings as a proxy for [DA] in health and PD, respectively. We therefore adjusted the parameters of GABA synapses in our model to be consistent with the experimental observation at high [DA], maintaining the maximal conductance ratio between D1 and D2 SPNs, and fitting the relaxation time of depression, individually for D1 and D2 SPNs (Fig. 1B).
- Ca^++^ dynamics: our current version of the model accounts for the stronger activation of calcium-dependent potassium channels in D2 SPNs compared to D1 SPNs, as reported experimentally (*13*) (Fig. 1C). This is mediated by a higher maximal conductance of the inward high-threshold calcium current in D2 SPNs.
- SPNs resonance: beta oscillations in the previous version of the model emerged from a rebound mechanism originated from an interaction between the intrinsic outward M-type potassium current and the network GABAergic synaptic inputs (*36*). In contrast, in this version of the model, SPN beta resonances (Fig. 2E) emerge from excitatory inputs and rely purely on the network GABAergic synaptic mechanism, as inferred from strengthened beta resonances following the absence of M-type input currents (Fig. S2 vs. Fig. 2E).
2. In support of biased competition under balanced input, the rate bias mechanism, and the coexistence of rate and coherence biases, are not fully consistent with reported rhythmic activity in PFC in the context of rule-based decisions:

- According to (*31*), modulated high beta oscillatory activity in PFC is associated with rule-based action selection, presumably mediated by preferential activation of relevant D1 SPNs (relevant GO pathway) vs. irrelevant D1 SPNs (irrelevant GO pathway) and D2 SPNs (NO-GO pathway), whereas alpha oscillatory activity in PFC is associated with inhibitory control, presumably mediated by preferential activation of D2 SPNs (NO-GO pathway) against D1 SPNs (GO pathway).
- Rate bias mechanism: only the slow integrator decoder is able to specifically read out rate biases.
  - For high beta inputs, we limit the analysis to compare the activity of relevant vs. irrelevant D1 SPNs for high input strength only, because this is the condition in which D1 SPN activity (GO pathway) surpasses the activity of D2 SPNs (NO-GO pathway) (Fig 2C). As seen in Figure 4A bottom, the slow integrator decoder is unable to reliably distinguish the activity of relevant and irrelevant D1 SPNs, hence it is incapable of responding according to the relevant feature of the stimulus. This is in contrast to the fast integrator decoder, which naturally reads out the coherence bias (Fig. 4A).
  - For alpha inputs, we limit the analysis to inputs of low strength only, because this is the condition in which D2 SPN activity surpasses the activity of D1 SPNs (Fig 2C). Even though the slow integrator decoder is able to respond according to the activity of D2 SPNs (Fig. S3A slow integrator panel), this is not due to the spectral content of the input, as was the case in (Fig. 4B), but is mediated by the low strength of the input (see Fig. 3A bottom). This is confirmed by identical dynamics under high beta oscillatory inputs (Fig. S3B slow integrator panels).
- Coexisting biases: the coexistence bias mechanism assumes inputs of low strength and a shift in the regime of operation of the read-out decoder (Fig. 3C), so that the decoder is able to respond either to the overall higher firing rate of D2 SPNs (slow integrator decoder), or to the higher coherence bias of D1 SPNs (fast integrator decoder).
  - The case of alpha inputs is the same here as in the rate bias mechanism. The slow integrator decoder indeed preferentially processes D2 SPNs activity bias, but this is not specific to a rhythmic input in the alpha frequency band (Fig. S3, slow integrator panels).
  - The dynamics under high beta oscillatory inputs of low strength is more difficult to predict, given the mismatch between the frequency of the cortical input at high beta frequency and SPNs resonance at low beta frequency (Fig. 2E). Are relevant D1 SPNs able to transform a high beta coherence bias in PFC input into a coherence bias with respect to irrelevant D1 SPNs and D2 SPNs? Figure S3B (fast integrator panels) shows that this is actually not the case: even though relevant D1 SPNs are more strongly synchronized than irrelevant D1 SPNs (Fig. S3B left vs. right fast integrator panels), D2 SPNs become synchronized more rapidly than D1 SPNs (Fig. S3B fast integrator panels), and even later on, D1 and D2 SPNs become similarly synchronized (Fig. S3B fast integrator panels).

### Figs S1 to S3

**Fig. S1.**
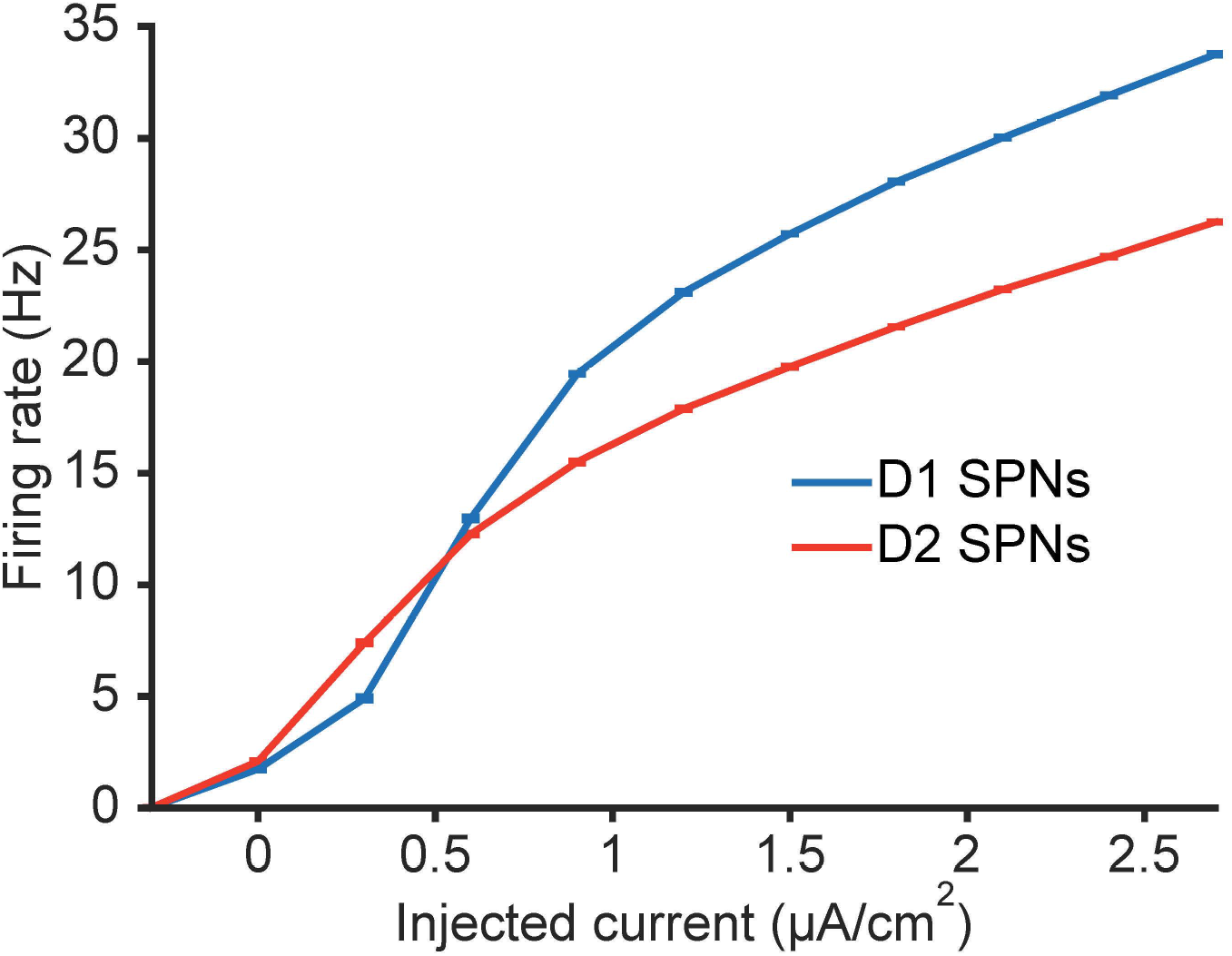
Averaged f-I curve of D1 and D2 SPNs under all-to-all connectivity. The curves represent the input-output relationship between injected current and time-averaged population firing rate under all-to-all connectivity. For this analysis, GABAergic conductances have been scaled down by the reported connectivity ratio (*11*), so that the effective conductance stays the same with respect to the conductance under asymmetric, sparse connectivity.

**Fig. S2.**
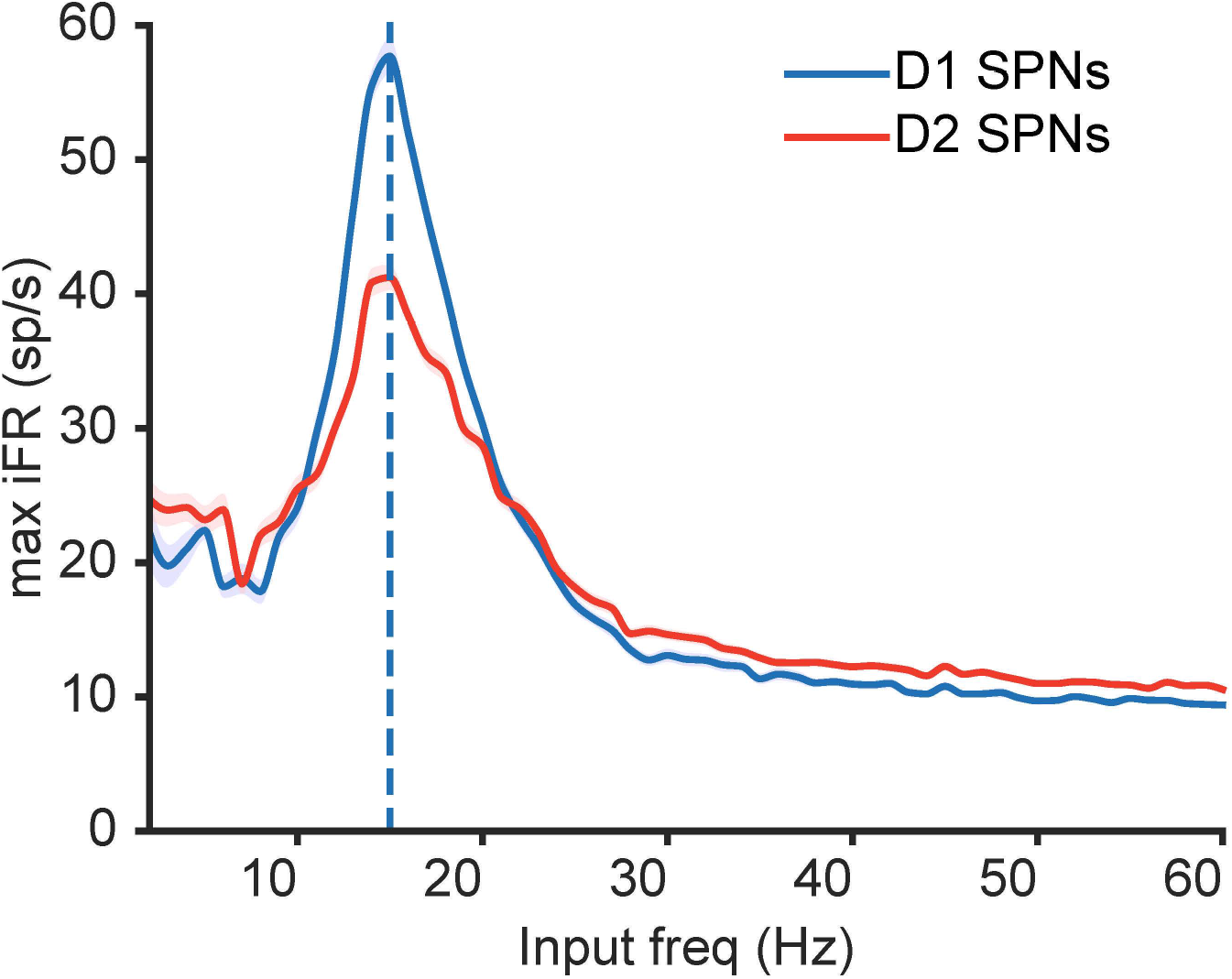
Resonance of SPNs for inputs of low strength in the absence of M-type input currents. The resonance is quantified in terms of maximum iFR, a measure of local population synchronization: across input frequencies (x-axis), the average of the peak iFR through all cycles is computed (y-axis).

**Fig. S3.**
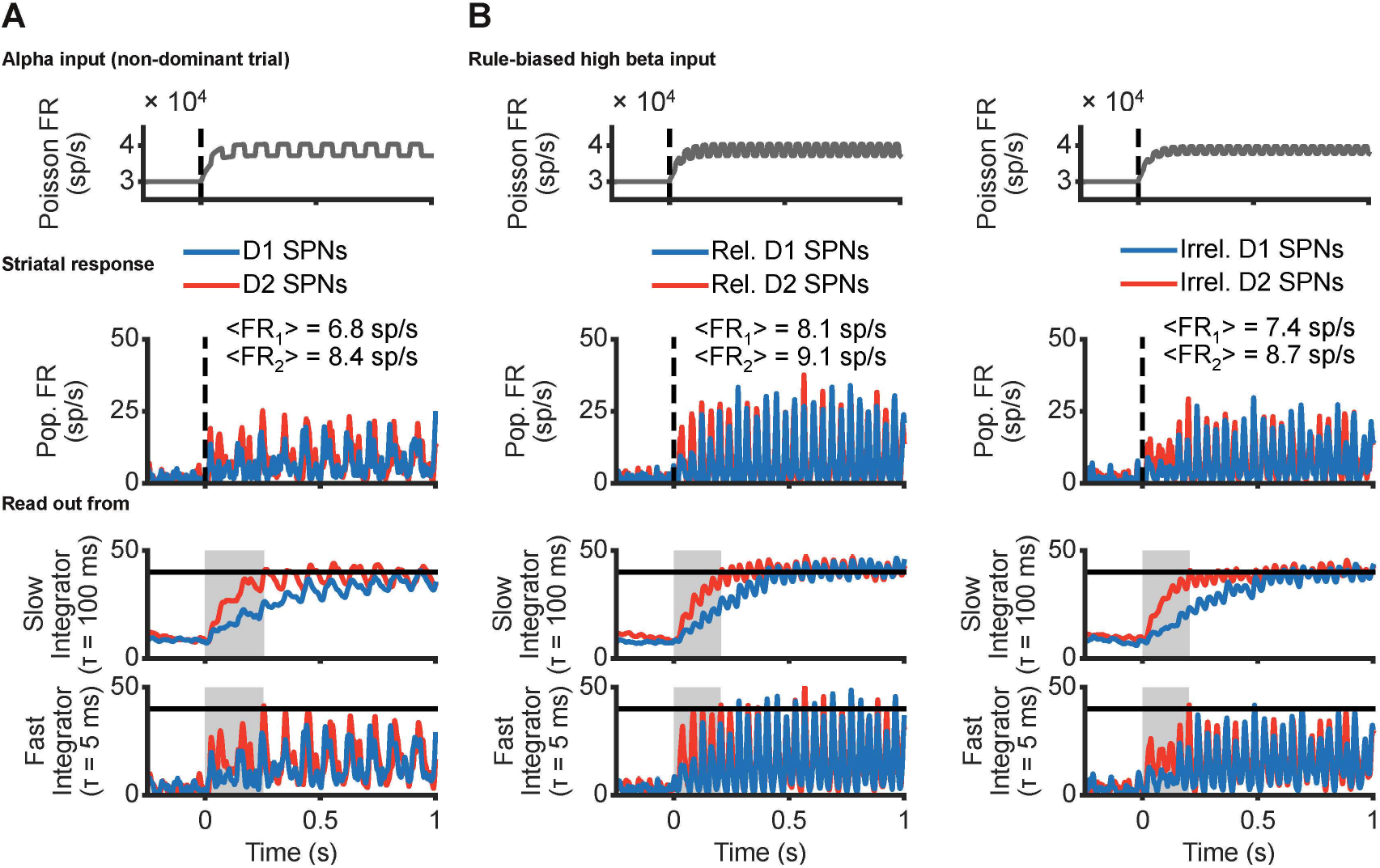
Striatal processing of low strength inputs oscillating at alpha and high beta frequencies. (**A**) Alpha input. Top: Low strength, alpha rhythmic input. Middle: Population FR varying in time (iFR). Bottom: Downstream readout of the activity of SPNs using distinct integration timescale. Response time is shaded in light gray. Response threshold at 40 sp/s (solid horizontal line). (**B**) Rule-biased high beta inputs (see main and supplementary text for details). Top: Relevant SPNs (left) receive more synchronized inputs (oscillations of higher amplitude) compared to irrelevant SPNs (right). Middle: Population FR varying in time (iFR). Bottom: Downstream readout of the activity of SPNs using distinct integration timescale. Response time is shaded in light gray. Response threshold at 40 sp/s (solid horizontal line).

### Materials and Methods

#### Model specification

Our striatal model consists of two populations of neurons, each population (*N* = 150) representing a distinct type of spiny projecting neurons (SPNs) according to dopamine receptor expression: D1R vs. D2R. Each SPN is modeled by a single compartment, representative of the soma. The membrane voltage *V* (mV) of each cell integrates input currents in time according to the following first order ordinary differential equation (ODE):

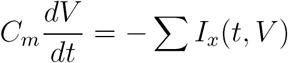

where *t* refers to time (ms), *C_m_* is the membrane capacitance (1*μ*F/cm^2^), and *I_x_* (*μ*A/cm^2^) spans three distinct input types: *I_intr_* for intrinsic input from ionic channels, *I_syn,int_* for synaptic inputs from striatal SPNs, and *I_syn,ext_* for synaptic inputs from external sources.

#### Intrinsic input currents

*I_intr_* consists of inward fast sodium *I_Na_*, outward fast delayed rectifier potassium *I_K_*, leak *I_L_*, outward M-type potassium *I_M_*, inward high-threshold calcium *I_Ca_*, and outward calcium-dependent potassium *I_KCa_* components. *I_Na_, I_K_, I_L_, I_M_*, and *I_Ca_* are modeled following the Hodgkin-Huxley formulation:

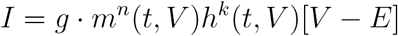

where *g* denotes the maximal conductance (mS/cm^2^), E is the reversal potential (mV), and *m*(*t, V*) and *h*(*t, V*) represent activation and inactivation gating variables of *n* and *k* order kinetics (*n, k* ≥ 0), respectively.

Ion channel gating variables, x in {m,h}, are described in time by a two-state kinetic first order ODE. They can be either expressed in terms of opening and closing channel rates, *α_x_*(V) and *β_x_*(V), as follows

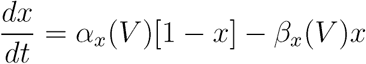

or in terms of the steady-state and the time constant, *x_∞_*(*V*) and *τ_x_*(*V*)

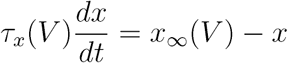

according to the relationship of *x*_∞_ and *τ_x_* with respect to *α_x_* and *β_x_*:

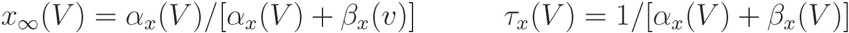

For the *I_Na_* current, *g_Na_* = 100*mS/cm*^2^, *E_Na_* = 50*mV*, *n_Na_* = 3 and *k_Na_* = 1. The rate functions (*α_Na_* and *β_Na_)* for the activation (*m_Na_*) and inactivation (*h_Na_*) gating variables are:

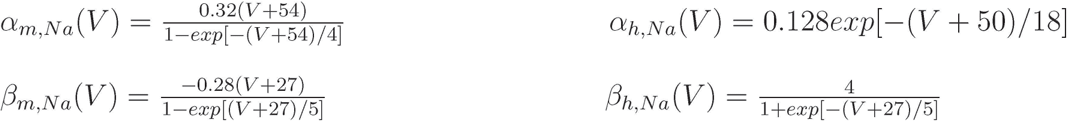

For the *I_K_* current, *g_K_* = 80*mS/cm*^2^, *E_K_* = –100*mV, n_K_* = 4 and *k_K_* = 0 (i.e., no inactivation). The rate functions (*α_K_* and *β*_K_) for the activation (*m_K_*) gating variable are:

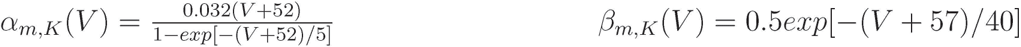

For the *I_L_* current, *E_K_* = –67mV, *n_K_* = 0 and *k_K_* = 0 (i.e., no gating variables). *g_L_* is slightly different between D1 and D2 SPNs: 0.097*mS/cm*^2^ and 0.1*mS/cm*^2^ respectively, which accounts for non-significantly different baseline activities between the two phenotypes (*39*).

For the *I_M_* current, *g_M_* = 1.3*mS/cm*^2^, *E_M_* = –100*mV*, *n_M_* = 1 and *k_M_* = 0 (i.e., no inactivation). The rate functions (*α_M_* and *β_M_*) for the activation (*m_M_*) gating variable are:

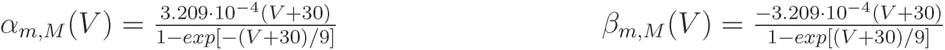

For the *I_Ca_* current, *E_Ca_* = 120*mV*, *n_Ca_* = 2 and *k_Ca_* = 0 (i.e., no inactivation). *g_Ca_* is different between D1 and D2 SPNs: 0.018*mS/cm*^2^ and 0.025*mS/cm*^2^ respectively, which accounts for faster and stronger activation of the *I_KCa_* current in D2 SPNs (13)(Fig. 1C). The rate functions (*α_Ca_* and *β_Ca_*) for the activation (*m_Ca_*) gating variable are:

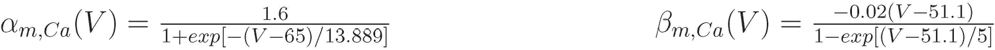

For the *I_KCa_, g_KCa_* = 0.2*mS/cm*^2^ and *E_KCa_* = –80*mV. I_KCa_*, however, activates by calcium concentration (in mM) instead of voltage (*n_KCa_* = 1 and *k_KCa_* = 0, i.e., no inactivation). For simplicity, the time constant of activation is considered fixed at _*τm,KCa*_([*Ca*]) = 120*ms*, whereas the steady-state of the activation (*m_KCa_*) gating variable is:

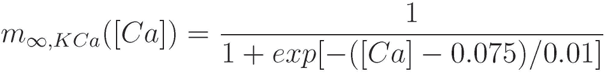

[*Ca*] varies according to the following calcium buffer dynamics

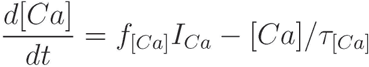

where 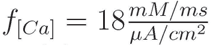 is the [*Ca*] accumulation factor and *τ*_[*Ca*]_ = 50*ms* represents the decay time in calcium concentration.

#### Synaptic input currents among SPNs

*I_syn,int_* refers to the inhibitory synaptic inputs between striatal SPNs mediated by GABAA receptors. For each SPN, *I_syn,int_* is modeled according to:

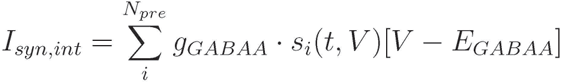

where *E_GABAA_* = –80*mV* represents the reversal potential and *g_gabaa_* refers to the maximal gabaergic conductance, whose value (0.65*mS/cm*^2^) is divisively normalized by the number of presynaptic cells of each phenotype (*N_pre_* = 150) and multiplicatively scaled by up to three factors. The first factor (0.635) applies specifically to presynaptic D2 SPNs to adjust for a higher first postsynaptic current emerging from D1 SPNs at high [DA] levels (*12*) (Fig. 1B). The second factor specifies the number of synaptic contacts between each pair of SPNs, which is randomly sampled according to the different proportion of synaptic connections (*11*) (Fig. 1A): 0.26 from D1 to D1 SPNs, 0.06 from D1 to D2 SPNs, 0.27 from D2 to D1 SPNs, 0.36 from D2 to D2 SPNs. The third factor (1.5) qualitatively accounts for stronger cross-inhibition (between D1 and D2 SPNs) compared to self-inhibition (within D1 and D2 SPNs) (*11*).

*s_i_*(*t, V*) represents presynaptic gating variables, whose dynamics evolves according to the following first order ODE:

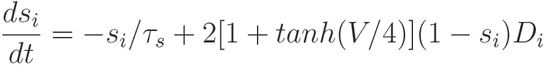

where the decay time of inhibition, *τ_s_*, is sampled from a normal distribution centered at 30.4ms with a standard deviation of 8.2ms to be in agreement with (*12*). *D_i_*(*t, V*) accounts for presynaptic short-term depression between SPN synapses:

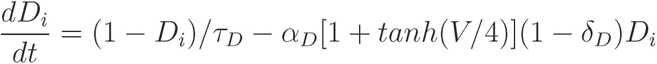

where *α_D_* = 2.305 and *δ_D_* = 0.35 control how much depression is accumulated after each spike, and *τ_D_* is the decay time of depression, which is significantly longer in D1 SPNs (1030*ms*) than in D2 SPNs (210ms). These parameters were adjusted according to (*12*)(Fig. 1B).

External synaptic input currents. *I_syn,ext_* refers to the long-range excitatory synaptic inputs to striatal SPNs from external sources, which are modeled according to stochastic Poisson spike trains. *I_syn,ext_* accounts for two distinct components: the background input, an asynchronous, uncorrelated, and unspecific input, whose strength determines the baseline activity of SPNs; and a task-dependent cortical input, assumed to be equal for D1 and D2 SPNs (*10*). For simplicity, both components are assumed to be mediated by AMPA receptors. Thus, for each SPN, *I_syn,ext_* is modeled according to:

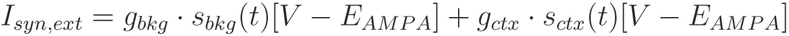

where *E_AM PA_* = 0*mV* represents the reversal potential. The maximal AMPA conductance is set to *g_x_* = 3.5*μS/cm*^2^ for both components *x* in {bkg, ctx}. *s_x_*(*t*) represents the gating variables, modeled as instantaneous jumps of magnitude 1 for each presynaptic spike followed by an exponential decay of *τ_s_* = 2*ms*. The two components differ, however, in terms of the Poisson process representing presynaptic firing.

For the background input, each SPN in each simulation is targeted by a different realization of a homogeneous Poisson process representing a total presynaptic firing of *τ_bkg_* = 30*kHz*, which can be mediated, for instance, by 15 thousand synapses (*40*) transmitting 2*sp/s* on average.

In contrast, the properties of the task-dependent cortical input differs in distinct simulations: the cortical input may be asynchronous (homogeneous Poisson process) or rhythmic (inhomogeneous Poisson process), it may be of high or low overall strength, and, if rhythmic, the input may oscillate at distinct frequency bands. To avoid sudden transitions at stimulus onset, the cortical input ramps up exponentially with a time constant of *τ_ctx_* = 40*ms*.

Specifically, the asynchronous input of high strength is mediated by a Poisson process representing a total presynaptic firing of 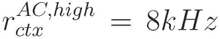. For high strength inputs, the amplitude of the rhythmic modulation is 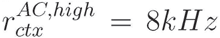. Low strength inputs scale down these values 5 times to 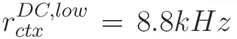 and 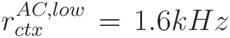. Also, in the context of rule-based decisions (*31*), the rhythmic modulation is scaled down 1.6 times in SPNs encoding irrelevant features with respect to SPNs encoding relevant features (Fig. 4A and S3B).

The frequency of the rhythmic, hence inhomogeneous, Poisson process is either centered at alpha frequencies *f_c_* = 10*Hz*, or at distinct beta bands: *f_c_* = 18Hz targets the resonance of both SPN types at low input strength, whereas *f_c_* = 25Hz and *f_c_* = 20*Hz* target, at high input strength, the resonance of D1 and D2 SPNs, respectively. All these frequencies represent the center around which the instantaneous frequency of each cycle jitters. In other words, the period of the oscillation slightly varies from cycle to cycle.

Cortical coordination is modeled as a smoothed square signal, arguably a better proxy of rhythmic synchronization in cortex compared to a sinusoid. The particular signal is implemented using periodic sigmoidal functions, of 1ms slope to generate smooth but sharp up and down transitions.

#### Data analysis

##### Population activity

Spiking multi-unit activity (MUA) is pooled over cells, separately for D1 and D2 SPNs (N = 150). The instantaneous firing rate is then estimated from MUA as follows: first, MUA is re-sampled from Δt = 0.05ms to Δt = 1ms, after which, the Nadaraya-Watson smoothing kernel regression algorithm is applied with a 5ms-bandwidth Epanechnikov kernel density estimator.

##### Read-out decoders

Downstream read-out of striatal output is estimated using two alternative decoders that accumulate SPNs activity at distinct pace (fast vs. slow integrator), but that are otherwise identical. The instantaneous firing rate output of the read-out decoder *r_dec_* evolves in time according to the following first order ODE:

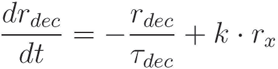

where *τ_dec_* is either 5*ms* or 100ms, respectively for the fast and slow integrator. *r_x_* represents the striatal output, i.e., SPNs instantaneous firing rate, where x is a label either referring to D1 or D2 SPNs. k is a scaling factor (in *s*^−1^) that is adjusted on a simulation basis so that the two read-out decoders are able to reach the same fixed response threshold at 40sp/s. *k_s1ow_* and *k_fast_* are set to {46,15.25, 40,14.7,14.7,14.7,14.2, 45, 45, 46} and {480,150, 200,110,110,110,122, 390, 380, 380} respectively, for the following ten conditions: low asynchronous input (Fig. 3A), high asynchronous input (Fig. 3B), low input resonance (Fig. 3C), high input D1 resonance (Fig. 3D), high input D2 resonance (Fig. 3E), rule-biased high beta input of high strength (Fig. 4A), alpha input of high strength (Fig. 4B), alpha input of low strength (Fig. S3A) and rule-biased high beta input of low strength, for relevant (Fig. S3B left) and irrelevant (Fig. S3B right) SPNs.

##### Implementation

The model was implemented in DynaSim (*41*), an open-source toolbox for simulating dynamical systems, compatible with MATLAB and GNU Octave. Simulations were executed using the fourth-order Runge-Kutta integration algorithm with a time step of Δ*t* = 0.05*ms*. The output of the read-out decoders was integrated with the variable time step ode45 built-in ODE solver in MATLAB/GNU Octave, which implements the fourth-order Dormand-Prince integration method.

